# N-terminal sumoylation of centromeric histone H3 variant Cse4 regulates its proteolysis to prevent mislocalization to non-centromeric chromatin

**DOI:** 10.1101/176206

**Authors:** Kentaro Ohkuni, Reuben Levy-Myers, Jack Warren, Wei-Chun Au, Yoshimitsu Takahashi, Richard E. Baker, Munira A. Basrai

**Affiliations:** Genetics Branch, Center for Cancer Research, National Cancer Institute, National Institute of Health, Bethesda, MD 20892; Biology Department, The College of William & Mary, Williamsburg, VA 23187; Department of Microbiology and Physiological Systems, University of Massachusetts Medical School, Worcester, MA 01655

**Keywords:** Sumoylation, Ubiquitination, Kinetochore, E3 ubiquitin ligase, Cse4, Slx5, Psh1, *Saccharomyces cerevisiae*

## Abstract

Stringent regulation of cellular levels of evolutionarily conserved centromeric histone H3 variant (CENP-A in humans, CID in flies, Cse4 in yeast) prevents its mislocalization to non-centromeric chromatin. Overexpression and mislocalization of CENP-A has been observed in cancers and leads to aneuploidy in yeast, flies, and human cells. Ubiquitin-mediated proteolysis of Cse4 by E3 ligases such as Psh1 and Sumo-Targeted Ubiquitin Ligase (STUbL) Slx5 prevent mislocalization of Cse4. Previously, we identified Siz1 and Siz2 as the major E3 ligases for sumoylation of Cse4. In this study, we identify lysine 65 (K65) in Cse4 as a SUMO site and show that sumoylation of Cse4 K65 regulates its ubiquitin-mediated proteolysis by Slx5. Strains expressing *cse4 K65R* exhibit reduced levels of sumoylated and ubiquitinated Cse4 *in vivo*. Furthermore, co-immunoprecipitation experiments reveal reduced interaction of cse4 K65R with Slx5. Defects in sumoylation of cse4 K65R contribute to increased stability and mislocalization of cse4 K65R under normal physiological conditions. Based on the increased stability of cse4 K65R in *psh1∆* strains but not in *slx5∆* strains, we conclude that Slx5 targets sumoylated Cse4 K65 for ubiquitination-mediated proteolysis independent of Psh1. In summary, we have identified and characterized the physiological role of Cse4 sumoylation and determined that sumoylation of Cse4 K65 regulates ubiquitin-mediated proteolysis and prevents mislocalization of Cse4 which is required for genome stability.

## Introduction

Evolutionarily conserved centromeric histone H3, CENP-A (Cse4 in *Saccharomyces cerevisiae*, Cnp1 in *Schizosaccharomyces pombe*, CID in *Drosophila*, and CENP-A in humans), is essential for faithful chromosome segregation. Overexpression of Cse4, CID, and CENP-A leads to its mislocalization to non-centromeric regions and contributes to aneuploidy in flies, yeast, and human cells, respectively (Heun et al. 2006; Moreno-Moreno et al. 2006; Collins et al. 2004; Au et al. 2008; Shrestha et al. 2017). Overexpression and mislocalization of CENP-A, which is observed in many cancers, is associated with poor prognosis and increased invasiveness (Tomonaga et al. 2003; Amato et al. 2009; Hu et al. 2010; Li et al. 2011; Wu et al. 2012; Lacoste et al. 2014; Athwal et al. 2015; Sun et al. 2016). Hence, characterization of pathways that regulate cellular levels of CENP-A and its homologs will help us to understand how overexpression and mislocalization of CENP-A contributes to tumorigenesis in human cells.

In budding yeast, ubiquitin-mediated proteolysis of Cse4 has been shown to regulate cellular levels of Cse4 and this prevents its mislocalization to non-centromeric regions (Collins et al. 2004). Proteolysis of Cse4 is regulated by several proteins: Psh1 (E3 ubiquitin ligase), Doa1 (WD-repeat protein), Fpr3 (proline isomerase), Ubp8 (ubiquitin protease), Rcy1 (F-box protein), and Ubr1 (E3 ubiquitin ligase) (Hewawasam et al. 2010; Ranjitkar et al. 2010; Au et al. 2013; Ohkuni et al. 2014; Canzonetta et al. 2015; Cheng et al. 2016; Cheng et al. 2017). Psh1-mediated proteolysis of Cse4 is regulated by its interaction with Spt16, a component of the FACT (facilitates chromatin transcription/transactions) complex, and casein kinase 2 (CK2) (Deyter and Biggins 2014; Hewawasam et al. 2014). Even though Psh1 interacts with the CENP-A targeting domain (CATD) and ubiquitinates lysine (K) residues (K131, K155, K163, and K172) in the C-terminus of Cse4 (Hewawasam et al. 2010; Ranjitkar et al. 2010), several observations support an important role of the N-terminus of Cse4 for regulating its ubiquitin-mediated proteolysis. For example, ubiquitination of cse4 ∆129 (a deletion of the entire N-terminus of Cse4) is barely detectable (Au et al. 2013). The stability of cse4 KC, where all K in the N-terminus of Cse4 are mutated to arginine, is at least three-fold higher than that of wild-type Cse4 (Au et al. 2013). Furthermore, the stability of cse4 KC is at least four-fold higher than that of cse4 CK, where all K in the C-terminus of Cse4 are mutated to arginine (Au et al. 2013). Proteolysis defects in N-terminus Cse4 mutants contribute to chromosome loss, as strains expressing *cse4 KC* exhibit a four-fold increase in chromosome segregation defects when compared to *cse4 CK* strains (Au et al. 2013). These results indicate that ubiquitination and interaction of Psh1 with the C-terminus of Cse4 is not sufficient for Cse4 ubiquitin-mediated proteolysis. Studies with fission yeast have also shown that the N-terminus of Cnp1 mediates ubiquitin-mediated proteolysis of Cnp1 to prevent its mislocalization to non centromeric chromatin (Gonzalez et al. 2014).

In addition to ubiquitination, we have recently reported that Cse4 is sumoylated by the small ubiquitin-related modifier (SUMO) E3 ligases Siz1 and Siz2 and that SUMO modification of Cse4 regulates its proteolysis (Ohkuni et al. 2016). We showed that *slx5*∆ strains exhibit defects in Cse4 proteolysis and localization that are independent of Psh1. Slx5 is a SUMO-targeted ubiquitin ligase which regulates many substrates with diverse biological functions such as transcription and maintaining genome stability. Slx5 associates with centromeric chromatin, and *slx5*∆ strains exhibit aneuploidy, spindle mispositioning, and aberrant spindle kinetics (van de Pasch et al. 2013). Given the multiplicity of Slx5 substrates, it is possible that the defect in Cse4 proteolysis in *slx5*∆ strains may be indirect due to defects in post-translational modifications of other substrates. Hence, we sought to identify sumoylation sites in the N-terminus of Cse4 and investigated how the interaction of Slx5 with sumoylated Cse4 regulates its proteolysis and localization. Here we report the identification of K65 in Cse4 as a SUMO site and show that Slx5-mediated ubiquitination of Cse4 K65 prevents its mislocalization to non-centromeric regions for genome stability.

## Materials and methods

### Yeast strains and plasmids

Table 1 and 2 describe the genotype of yeast strains and plasmids used for this study, respectively. Yeast cells were grown in rich (YPD: 1 % yeast extract, 2 % bacto-peptone, 2 % glucose) or synthetic complete (SC) media containing 2 % glucose, or 2 % raffinose, or 2 % raffinose + 2 % galactose.

**Table 1.** *S. cerevisiae* stains used in this study.

| Strains | Parent | Genotype | Reference |
| --- | --- | --- | --- |
| BY4741 |  | *MATa his3Δ1 leu2Δ0 met15Δ0 ura3Δ0* | Open Biosystems |
| BY4742 |  | *MATα his3Δ1 leu2Δ0 lys2Δ0 ura3Δ0* | Open Biosystems |
| YMB9470 | BY4741/BY4742 | *MATα cse4Δ::kanMX6 [pRS415-6His-3HA-CSE4]* | This study |
| YMB9471 | BY4741/BY4742 | *MATα cse4Δ::kanMX6 psh1Δ::kanMX6 [pRS415-6His-3HA-CSE4]* | This study |
| YMB9472 | BY4741/BY4742 | *MATα cse4Δ::kanMX6 slx5Δ::kanMX6 [pRS415-6His-3HA-CSE4]* | This study |
| YMB9547 | BY4741/BY4742 | *MATα cse4Δ::kanMX6 [pRS415-6His-3HA-cse4 K65R]* | This study |
| YMB9548 | BY4741/BY4742 | *MATα cse4Δ::kanMX6 psh1Δ::kanMX6 [pRS415-6His-3HA-cse4 K65R]* | This study |
| YMB9549 | BY4741/BY4742 | *MATα cse4Δ::kanMX6 slx5Δ::kanMX6 [pRS415-6His-3HA-cse4 K65R]* | This study |
| YMB9694 | BY4741 | *MATa SLX5-Myc:kanMX6* | This study |
| YPH1018 |  | *MATα ura3-52 lys2-801 ade2-101 trp1Δ63 his3Δ200 leu2Δ1 CFIII (CEN3L.YPH278) HIS3 SUP11* | P. Hieter |

**Table 2.** Plasmids used in this study.

| Plasmid | Relevant characteristics | Reference |
| --- | --- | --- |
| pYES2 | Vector (*pGAL*, $2\mu$, *URA3*) | (Boeckmann et al. 2013) |
| pMB433 | Vector (p426-*pGAL1*, $2\mu$, *URA3*) | (Mumberg et al. 1994) |
| pMB1344 | pYES2-*pGAL-8His-HA-cse4 16KR* ($2\mu$, *URA3*) | (Ohkuni et al. 2016) |
| pMB1345 | pYES2-*pGAL-8His-HA-CSE4* ($2\mu$, *URA3*) | (Ohkuni et al. 2016) |
| pMB1458 | pMB433-*pGAL-6His-3HA-CSE4* ($2\mu$, *URA3*) | (Au et al. 2013) |
| pMB1597 | pMB433-*pGAL-3HA-CSE4* ($2\mu$, *URA3*) | (Ohkuni et al. 2016) |
| pMB1725 | pRS415-*6His-3HA-CSE4* (*CEN*, *LEU2*) | This study |
| pMB1749 | pRS415-*6His-3HA-cse4 K65R* (*CEN*, *LEU2*) | This study |
| pMB1762 | pYES2-*pGAL-8His-HA-cse4 K65R* ($2\mu$, *URA3*) | This study |
| pMB1791 | pMB433-*pGAL-6His-3HA-cse4 K65R* ($2\mu$, *URA3*) | This study |
| pMB1814 | pMB433-*pGAL-3HA-cse4 K65R* ($2\mu$, *URA3*) | This study |

### Cell lysate preparation for pull down assays and coimmunoprecipitation (co-IP)

For pull down assays and immunoprecipitation, cell lysates were prepared as follows. 50 OD_600_ of logarithmically growing cells were pelleted, rinsed with sterile water, and suspended in 0.5 ml of guanidine buffer (0.1 M Tris-HCl at pH 8.0, 6.0 M guanidine chloride, 0.5 M NaCl) for sumoylation assay, or Ub lysis buffer (20 mM Na_2_HPO_4_, 20 mM NaH_2_PO_4_, 50 mM NaF, 5 mM tetra-sodium pyrophosphate, 10 mM beta-glycerophosphate, 2 mM EDTA, 1 mM DTT, 1 % NP-40, 1 mM phenylmethylsulfonyl fluoride (PMSF), 5 mM N-ethylmaleimide (NEM), 1x protease inhibitor cocktail (Sigma, P8215)) for ubiquitin pull down assay, or IP lysis buffer (50 mM Tris-HCl at pH 8.0, 5 mM EDTA, 1 % TritonX-100, 150 mM NaCl, 50 mM NaF, 10 mM β-glycerophosphate, 1 mM PMSF, 1x protease inhibitor cocktail) for coimmunoprecipitation. Cells were homogenized with 0.5 mm diameter glass beads or Matrix C (MP Biomedicals) using a bead beater (MP Biomedicals, FastPrep-24 5G). Cell lysates were clarified by centrifugation at 6,000 rpm for 5 min and protein concentration was determined using a DC protein assay kit (Bio-Rad). Samples containing equal amounts of protein were brought to a total volume of 1 ml with appropriate buffer.

### Sumoylation assay

*In vivo* sumoylation was assayed in crude yeast extracts using Ni-NTA agarose beads to pull down His-HA-tagged Cse4 as described previously (Ohkuni et al. 2015). Cell lysates were incubated with 100 μl of Ni-NTA superflow beads (Qiagen, 30430) overnight at 4 °C. After being washed with guanidine buffer one time and with breaking buffer (0.1 M Tris-HCl at pH 8.0, 20 % glycerol, 1 mM PMSF) four times, beads were incubated with 2x Laemmli buffer including imidazole at 100°C for 10 min. The protein samples were analyzed by SDS-PAGE and Western Blotting.

### Ubiquitin pull down assay

Ubiquitin-pull down assay was performed as described previously (Au et al. 2013). Cell lysates were incubated with 25 μl of tandem ubiquitin-binding entities Agarose-TUBE1 (LifeSensors, Inc. UM401) overnight at 4 °C. After being washed with TBS-T (20 mM Tris-HCl at pH 7.5, 150 mM NaCl, 0.1 % Tween20) three times, beads were incubated with 2x Laemmli buffer at 100°C for 5 min. The protein samples were analyzed by SDS-PAGE and Western Blotting.

### Coimmunoprecipitation (Co-IP)

Cell lysates were incubated with 25 μl of Anti-HA-Agarose antibody (Sigma, A2095) overnight at 4 °C. After being washed with IP lysis buffer three times, beads were incubated with 2x Laemmli buffer at 100°C for 5 min. The protein samples were analyzed by SDS-PAGE and Western Blotting.

### Protein stability assay

Protein stability assays were performed as described previously (Ohkuni et al. 2016). Logarithmically growing cells in YPD were treated with cycloheximide (CHX) (20 μg/ml). Protein extracts were prepared from cells at the indicated time points and levels of Cse4 were quantitated by western blot analysis. Multiple independent measurements were made, and means are given on the applicable figures ± the standard error of the mean with the number of replicates given in parentheses. Statistical significance was assessed by one-way ANOVA (P = 0.0075) with selected pairwise post hoc tests. Bonferroni correction was applied to adjust P values for multiple comparisons.

### SDS-PAGE and Western Blotting

SDS-PAGE was performed using 4-12 % Bis-Tris gels (Novex). After running the gels, proteins were transferred onto nitrocellulose membrane (VitaScientific, DBOC80014). Western blotting was performed according to a standard protocol. SuperSignal West Pico Chemiluminescent Substrate (Thermo Scientific, 34078), or SuperSignal West Pico PLUS Chemiluminescent Substrate (Thermo Scientific, 34580), or One Component Chemiluminescent Substrate (Rockland, UniGlow-0100) was used for the detection. Protein levels were quantified using Gene Tools software (version 3.8.8.0) from SynGene (Frederick, MD) or Image Lab software (version 6.0.0) from Bio-Rad Laboratories, Inc (Hercules, CA). Antibodies were as follows: anti-HA (12CA5) mouse (Roche, 11583816001), anti-HA rabbit (Sigma, H6908), anti-c-Myc (A-14) (Santa Cruz Biotechnology, sc-789), anti-Smt3 (y-84) (Santa Cruz Biotechnology, sc-28649), anti-Tub2 antibodies (Basrai laboratory), and anti-FLAG (Sigma, F3165). Anti-HA was used at a dilution from 1:1000 to 1:5000, and anti-c-Myc, anti-Smt3, anti-FLAG, and anti-Tub2 were used at a dilution from 1:3000 to 1:5000. Secondary antibodies were ECL Mouse IgG, HRP-Linked Whole Ab (GE Healthcare Life Sciences, NA931V) or ECL Rabbit IgG, HRP-linked whole Ab (GE Healthcare Life Sciences, NA934V) at 1:5000 dilution.

### Chromosome spreads

Chromosome spreads were performed as described previously (Crotti and Basrai 2004; Collins et al. 2004) with some modifications. Cells were grown in YPD logarithmically. 16B12 Mouse anti-HA antibody (Covance, Babco; MMS-101P) was used as primary antibody at 1:2500 dilution. Cy3 conjugated Goat anti mouse (Jackson ImmunoResearch Laboratories, Inc., 115165003) was used as secondary antibody at 1:5000 dilution. DNA was visualized by DAPI (4’,6-diamidino-2-phenylindole) staining (1 μg/ml in phosphate-buffered saline) mounted in antifade mountant (Molecular Probes, P36935). Cells were examined under an Axioskop 2 (Zeiss) fluorescence microscope equipped with a Plan-APOCHROMAT 100X (Zeiss) oil immersion lens. Image acquisition and processing were performed with the IP Lab version 3.9.9 r3 software (Scanalytics, Inc.).

### Chromosome loss assay

Chromosome loss assays were performed as described previously (Spencer et al. 1990; Au et al. 2008). Briefly, strains containing a nonessential reporter chromosome were plated on synthetic defined medium with limiting adenine plus 2 % galactose/raffinose and incubated at 30°C for 5 days. Loss of the reporter chromosome results in red sectors in an otherwise a white colony. Colonies that are at least half red indicate loss of the reporter chromosome in the first cell division. At least 2,500 colonies were assayed from three individual transformants for each strain.

## Results

### Cse4 K65 is a target of sumoylation

The consensus motif Ψ-K-x-D/E (Ψ is a hydrophobic residue, K is the lysine to be conjugated to SUMO, x is any amino acid, D or E is an acidic residue) is frequently found in sumoylated substrates (Sampson et al. 2001; Rodriguez et al. 2001; Bernier-Villamor et al. 2002). We identified lysine (K) 65 (64-S**K**SD-67) in the N-terminus of Cse4 and K215/K216 (214-M**KK**D-217) in the C-terminus of Cse4 as potential sumoylation sites. Given the importance of the Cse4 N-terminus in ubiquitination and proteolysis (Au et al. 2013), we examined whether Cse4 K65 is sumoylated *in vivo* and further characterized its role in regulating proteolysis and localization of Cse4. We assayed the sumoylation status of transiently overexpressed *8His-Hemagglutinin (HA)-CSE4*, *8His-HA-cse4 16KR* (all K mutated to R), or *8His-HA-cse4 K65R* (K65 mutated to R65). Pull down experiments were done using Ni-NTA agarose beads, and SUMO-modified Cse4 species were detected using anti-Smt3 antibody (Figure 1). At least three distinct sumoylated Cse4 species were observed in wild-type cells (Figure 1A, denoted by arrows), as previously described (Ohkuni et al. 2016). These SUMO-modified species were not detected in strains expressing vector alone or *8His-HA-cse4 16KR*. Strains expressing *8His-HA-cse4 K65R* showed greatly reduced levels of Cse4 sumoylation even though similar levels of Cse4 and cse4 K65R were pulled down (Figure 1, A and B). Hence, we conclude that Cse4 K65 is sumoylated in wild-type cells.

**Figure 1.**
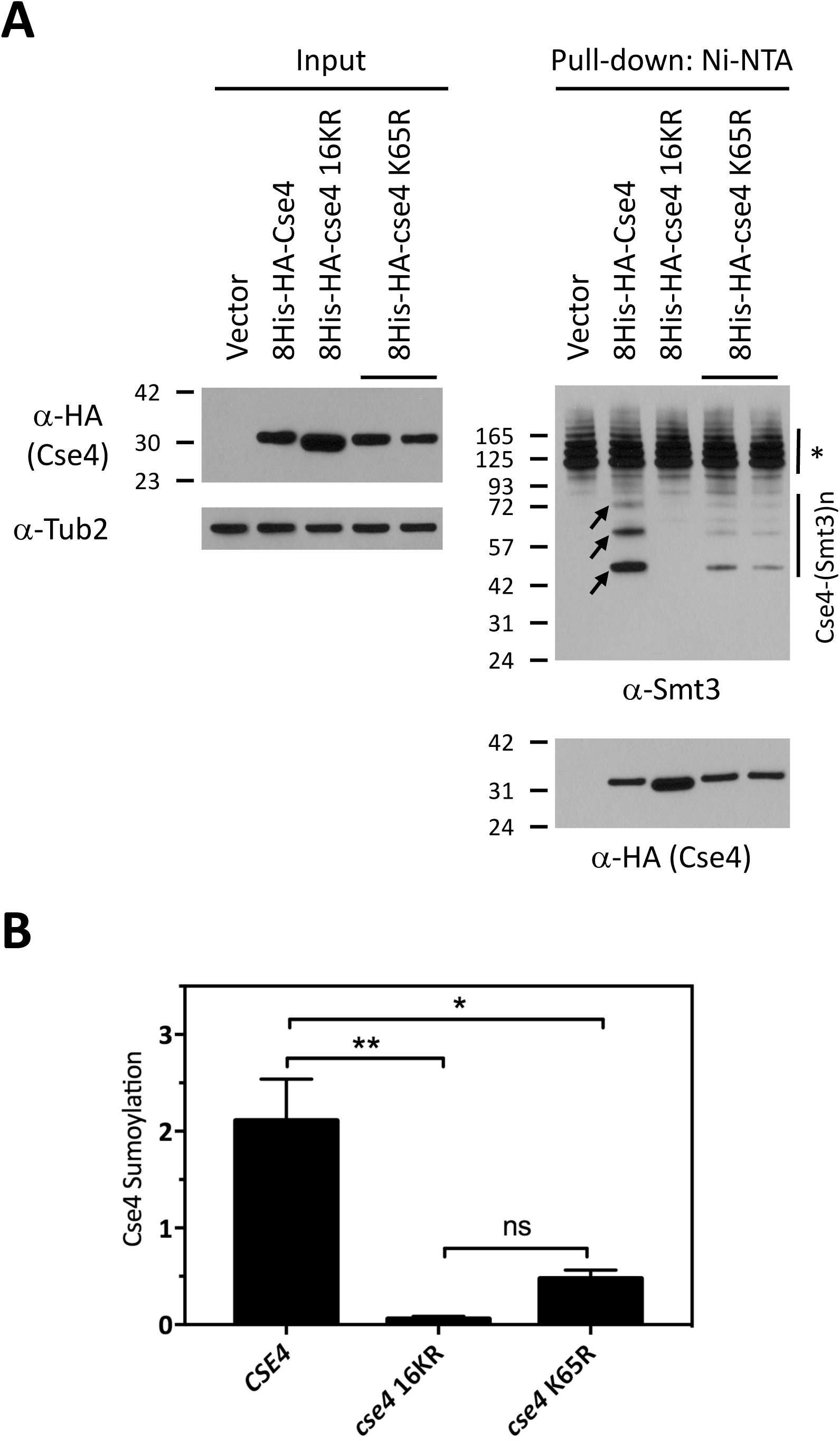
Cse4 K65 is sumoylated *in vivo*. (A) Wild-type strain (BY4741) transformed with vector (pYES2), *pGAL-8His-HA-CSE4* (pMB1345), *pGAL-8His-HA-cse4 16KR* (pMB1344), or *pGAL-8His-HA-cse4 K65R* (pMB1762) was grown in raffinose/galactose (2%) for 4 hours to induce expression of Cse4. Sumoylation of 8His-HA-Cse4 and nonmodified 8His-HA-Cse4 was assayed using protein extracts after pull down on Ni-NTA beads followed by western blot analysis with anti-Smt3 and anti-HA (Cse4) antibodies, respectively. At least three high molecular weight bands of 8His-HA-Cse4 (arrows) were detected. Input samples were analyzed using anti-HA (Cse4) and anti-Tub2 antibodies served as loading control for input. Asterisk shows nonspecific sumoylated proteins that bind to beads. (B) Quantitation of sumoylation in arbitrary density units (normalized to non-modified Cse4 probed by anti-HA in pull down sample) determined in multiple experiments (N = 3, 3, 2 for wild-type, Cse4 16KR, and Cse4 K65R, respectively). Statistical significance was assessed by one-way ANOVA (P = 0.007) followed by all possible pairwise comparisons of the means with Bonferroni correction. **, P < 0.01; *, P < 0.05; ns, not significant.

### Cse4 ubiquitination is reduced in a *cse4 K65R* mutant

The reduced sumoylation of cse4 K65R prompted us to investigate if sumoylation of Cse4 K65 affects the ubiquitination of Cse4. We assayed the level of ubiquitinated Cse4 by performing an affinity pull down assay using agarose with tandem ubiquitin-binding entities (Ub^+^) from strains transiently overexpressed either *3HA-CSE4* or *3HA-cse4 K65R* (Figure 2). A faster migrating species (Figure 2A, asterisk) was observed in all strains expressing Cse4. These faster migrating Cse4 species were similar in size to that in the input lanes. These species were also observed in experiments with *cse4 16KR* mutant (Au et al. 2013). Since cse4 16KR cannot be ubiquitinated, this faster migrating species represents unmodified Cse4, which likely interacts with ubiquitinated proteins such as canonical histones. Ubiquitinated Cse4 was detected as a laddering pattern in wild-type cells expressing *3HA-CSE4* but was absent in strains with vector alone (Figure 2A). However, reduced levels of ubiquitination were observed in strains expressing *3HA-cse4 K65R* (Figure 2, A and B). The lack of a drastic reduction in the levels of ubiquitinated cse4 K65R is not surprising given that the other fifteen lysines can still be ubiquitinated. These results show that Cse4 K65 contributes to Cse4 ubiquitination.

**Figure 2.**
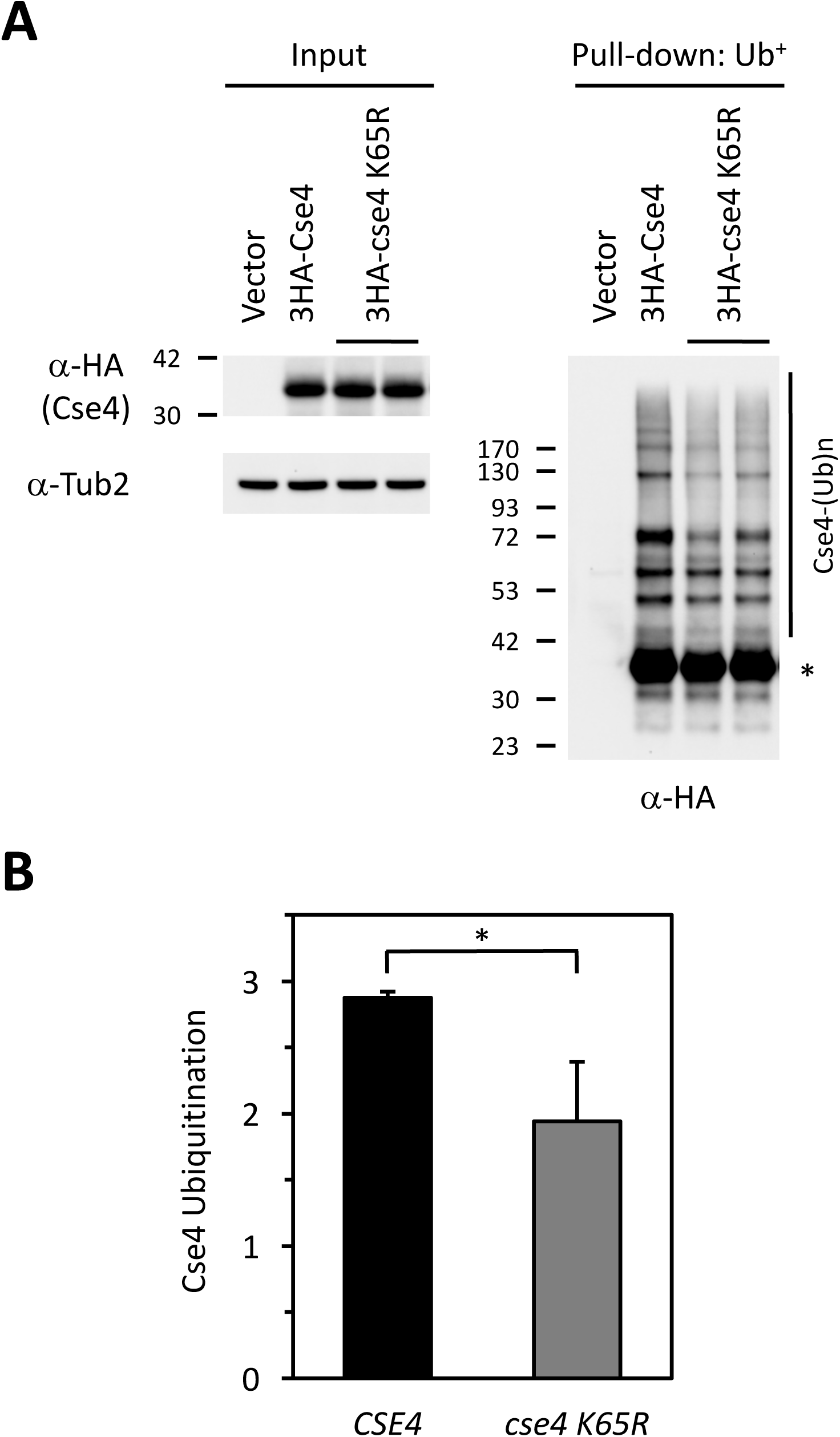
Cse4 ubiquitination is reduced in a *cse4 K65R* mutant. (A) Protein extracts were prepared from wild-type strain (BY4741) transformed with vector (pMB433), *pGAL-3HA-CSE4* (pMB1458), or *pGAL-3HA-cse4 K65R* (pMB1791) grown in raffinose/galactose (2%) for 4 hours to induce expression of Cse4. Agarose-TUBE1 was used to pull down tandem ubiquitin binding entities and ubiquitination levels of Cse4 were detected by western blot analysis with anti-HA antibody. Input samples were analyzed using anti-HA (Cse4) and anti-Tub2 antibodies. Asterisk shows non-modified Cse4. (B) Quantitation of ubiquitination in arbitrary density units (normalized to levels of Cse4 in input) in replicate determinations (N = 3, 4 for HA-Cse4 and HA-Cse4 K65R, respectively). Statistical significance was assessed by unpaired t test (*, P < 0.05).

### Cse4 K65 is regulated by STUbL Slx5

We have previously shown that STUbL Slx5 interacts with Cse4 and promotes its ubiquitin-mediated proteolysis (Ohkuni et al. 2016). Slx5 contains SUMO-interacting motifs (SIMs), which interact with sumoylated substrates (Xie et al. 2007; Xie et al. 2010). We reasoned that reduced sumoylation and ubiquitination of cse4 K65R may be due to reduced interaction of cse4 K65R with Slx5. Co-immunoprecipitation (Co-IP) experiments were done using strains co-expressing either *3HA-CSE4* or *3HA-cse4 K65R* with *SLX5-Myc* (Figure 3). Consistent with our previous results (Ohkuni et al. 2016), immunoprecipitation with anti-HA showed an interaction between wild-type Cse4 and Slx5 (Figure 3A). In contrast, reduced interaction was observed between cse4 K65R and Slx5 in two independent strains (Figure 3, A and B). Hence, we conclude that sumoylation of Cse4 K65 contributes to its interaction with Slx5.

**Figure 3.**
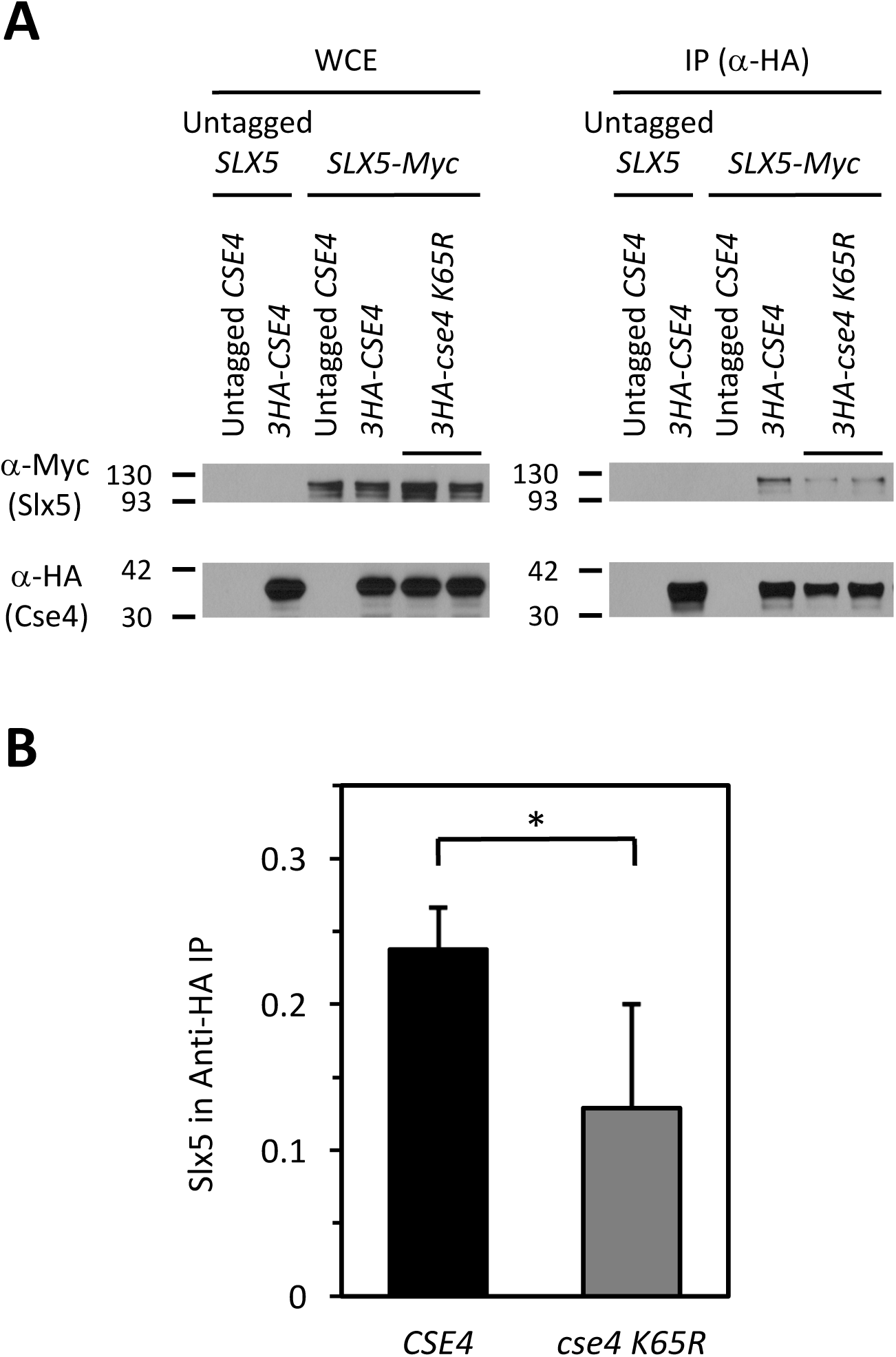
K65R mutation in Cse4 contributes to reduced interaction with Slx5. (A) Protein extracts were prepared from the indicated strains grown in raffinose/galactose (2%) for 4 hours to induce expression of Cse4 and incubated with anti-HA agarose antibody. Samples were resolved by SDS-PAGE and levels of Slx5 and Cse4 were detected by western blot analysis with anti-Myc and anti-HA antibodies, respectively. Isogenic yeast strains were: untagged (BY4741), or Slx5-Myc (YMB9694) strain transformed with vector (pMB433), *pGAL-3HA-CSE4* (pMB1458), or *pGAL-3HA-cse4 K65R* (pMB1791). (B) Levels of Slx5 (α-Myc, IP) after immunoprecipitation of 3HA-Cse4 were quantified in arbitrary density units after normalization to Cse4 levels in the IP (α-HA, IP) in three independent experiments (N = 3, 5 for HA-Cse4 and HA-Cse4 K65R, respectively). Statistical significance of the normalized values was assessed by unpaired t test (*, P < 0.05).

### Sumoylation of Cse4 K65 regulates its proteolysis and localization under normal physiological conditions

To determine if defects in sumoylation and ubiquitination of cse4 K65R contribute to a defect in proteolysis of cse4 K65R, we performed protein stability assays using cells expressing either *CSE4* or *cse4 K65R* from its endogenous promoter. Cells were grown logarithmically and then treated with cycloheximide (CHX) to inhibit protein translation. Whole cell extracts were assayed for Cse4 levels at different times after CHX treatment. Cse4 was rapidly degraded (t_1/2_ = 33.5 ± 0.8 min) in wild-type strains, whereas cse4 K65R was modestly stabilized (t_1/2_ = 50.8 ± 6.1 min) (Figure 4, A and B). These observations indicate that K65 of Cse4 regulates its proteolysis.

**Figure 4.**
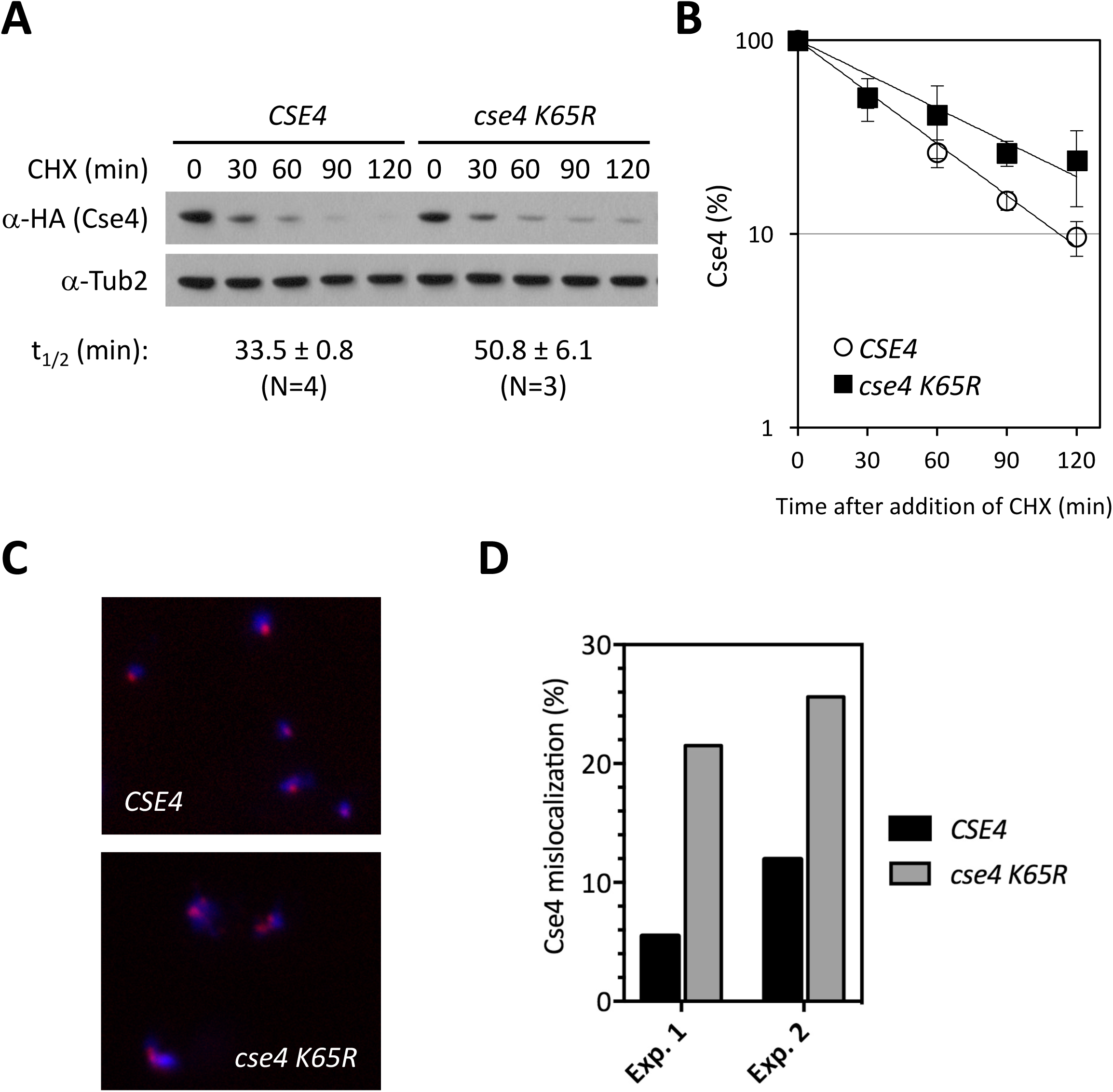
Sumoylation of Cse4 K65 regulates its proteolysis and localization under normal physiological conditions. (A) Protein extracts from *CSE4* (YMB9470) or *cse4 K65R* (YMB9547) strain were prepared using logarithmically growing cells, treated with CHX (20 μg/ml) for various time points. Blots were probed with anti-HA (Cse4) or anti-Tub2 (loading control) antibody. A representative blot is shown. Cse4 protein half-life (t_1/2_) is reported as the mean ± SEM of multiple biological repeats. The difference in t_1/2_ is statistically significant (P = 0.013). See materials and methods for details. (B) Kinetics of turnover. The graph shows the percentage of Cse4 signals normalized to Tub2 at the indicated time points in wild-type and *cse4 K65R* mutant. Error bars indicate standard deviations of the means. (C) Chromosome spreads of logarithmically-grown wild-type (YMB9470) or *cse4 K65R* (YMB9547) cells. DAPI (blue) and α-HA (red) staining were used to visualize DNA and Cse4 localization, respectively. In wild-type strains, Cse4 is predominantly localized to one to two kinetochore clusters. In mutant strains, mislocalization of Cse4 is observed as multiple foci or diffused localization throughout the nucleus. (D) Quantification of Cse4 mislocalization in two independent experiments. The increased mislocalization observed for the *cse4 K65R* strain was highly significant (P < 0.001) in both biological replicates (Chi-square test, N = 237, 237 and N = 251, 281 for wild-type and *cse4 K65R*, respectively in replicate 1 and replicate 2).

Previous studies have shown that defects in ubiquitin-mediated proteolysis of Cse4 in *slx5*∆ and *psh1∆* strains lead to mislocalization of Cse4 to non-centromeric regions under normal physiological conditions (Ohkuni et al. 2016). Therefore, we examined whether the increased stability observed for *cse4 K65R* expressed from its own promoter leads to mislocalization to non-centromeric regions. Chromosome spreads were used to visualize localization of chromatin-bound Cse4. In wild-type cells, Cse4 localization is usually restricted to one or two foci, corresponding to kinetochore clusters (Figure 4, C and D). In contrast, a higher incidence of multiple or diffuse cse4 K65R foci were observed under the same condition (Figure 4, C and D). We conclude that ubiquitin-mediated proteolysis of Cse4 via sumoylation of K65 prevents its mislocalization to non-centromeric chromatin.

### SUMO modification of Cse4 K65 is required for Slx5-mediated proteolysis independent of Psh1

Based on the reduced interaction of cse4 K65R with Slx5 (Figure 3), we predicted that the stability of cse4 K65R would not differ significantly from that of Cse4 in *slx5∆* strains under normal physiological conditions. In agreement with previous results (Ohkuni et al. 2016), protein stability assays showed that Cse4 was more stable in the *slx5*∆ strain (t_1/2_= 43.0 ± 6.5 min) when compared to the wild-type strain (t_1/2_= 33.5 ± 0.8 min) (Figure 5, A and B). Consistent with our hypothesis, the stability of cse4 K65R (t_1/2_= 43.5 ± 2.7 min) was not significantly different than that of Cse4 (t_1/2_ = 43.0 ± 6.5 min) in the *slx5*∆ strain (Figure 5, A and B).

**Figure 5.**
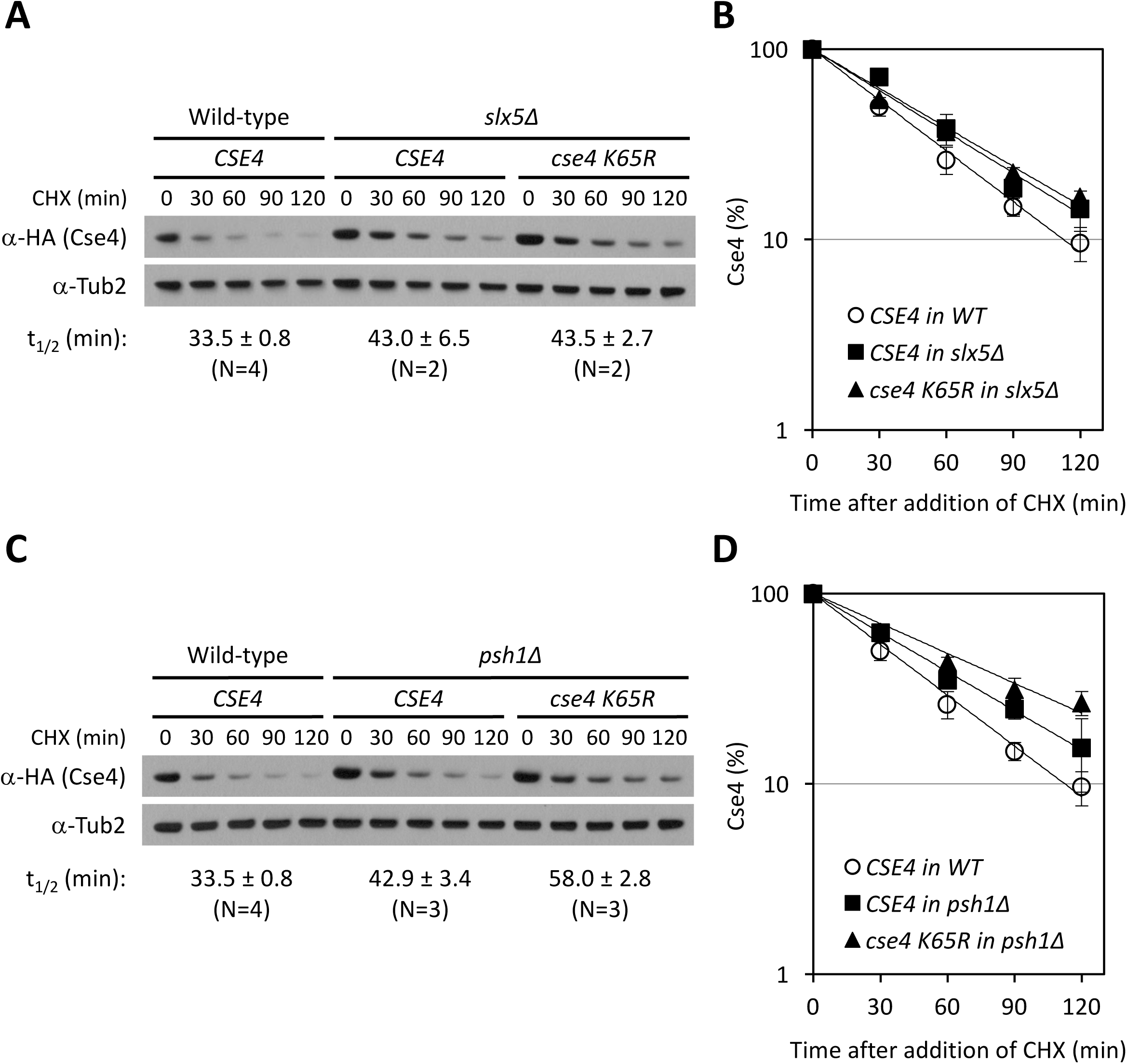
Cse4 K65 sumoylation regulates its proteolysis by Slx5 in a Psh1-independent manner. (A and C) Protein extracts were prepared using logarithmically growing cells, treated with CHX (20 μg/ml) for various time points. Blots were probed with anti-HA (Cse4) or anti-Tub2 (loading control) antibody. A representative blot is shown. Cse4 protein half-lives (t_1/2_) are reported as the mean ± SEM of multiple biological repeats. The difference in t_1/2_ between wild-type Cse4 and Cse4 K65R is statistically significant in the *psh1*∆ strain (P = 0.043) but not in the *slx5*∆ strain (P > 0.99). See materials and methods for details. (B and D) Kinetics of turnover. The graph shows the percentage of Cse4 signals normalized to Tub2 at the indicated time points in the indicated strains. Error bars indicate standard deviations of the means. (A) Strains analyzed were YMB9470 (*CSE4*), YMB9472 (*CSE4 slx5*∆), and YMB9549 (*cse4 K65R slx5*∆). (C) Strains analyzed were YMB9470 (*CSE4*), YMB9471 (*CSE4 psh1*∆), and YMB9548 (*cse4 K65R psh1*∆).

We next analyzed the stability of Cse4 using a *psh1*∆ strain. Protein stabililty assays showed that Cse4 was more stable in a *psh1*∆ strain (t_1/2_= 42.9 ± 3.4 min) compared to that observed in a wild type strain (t_1/2_= 33.5 ± 0.8 min) under normal physiological conditions (Figure 5, C and D). The stability of cse4 K65R (t_1/2_ = 58.0 ± 2.8 min) was much higher than that of Cse4 in the *psh1∆* strain (t_1/2_ = 42.9 ± 3.4 min) (Figure 5, C and D). Hence, we conclude that SUMO modification of Cse4 K65 is required for Slx5-mediated proteolysis that is independent of Psh1.

### Sumoylation of Cse4 K65 regulates faithful chromosome segregation

To determine the physiological consequence of mislocalized cse4 K65R on genome stability, we used a colony color assay to measure the loss of a nonessential reporter chromosome. Previous studies have shown that strains overexpressing *cse4 16KR* exhibit increased chromosome loss compared to *GAL-CSE4* (Au et al. 2008). However, no measurable increase in chromosome loss was observed in cells expressing *cse4 16KR* from its native promoter (Au et al. 2008). Therefore, we measured chromosome loss using wild type strain transformed with vector alone, *GAL-CSE4* and *GAL-cse4 K65R*. The *GAL-cse4 K65R* strain exhibits 1.6-fold higher chromosome loss compared to *GAL-CSE4* strains (Figure 6, P < 0.001). These results show that sumoylation of cse4 K65 regulates faithful chromosome segregation.

**Figure 6.**
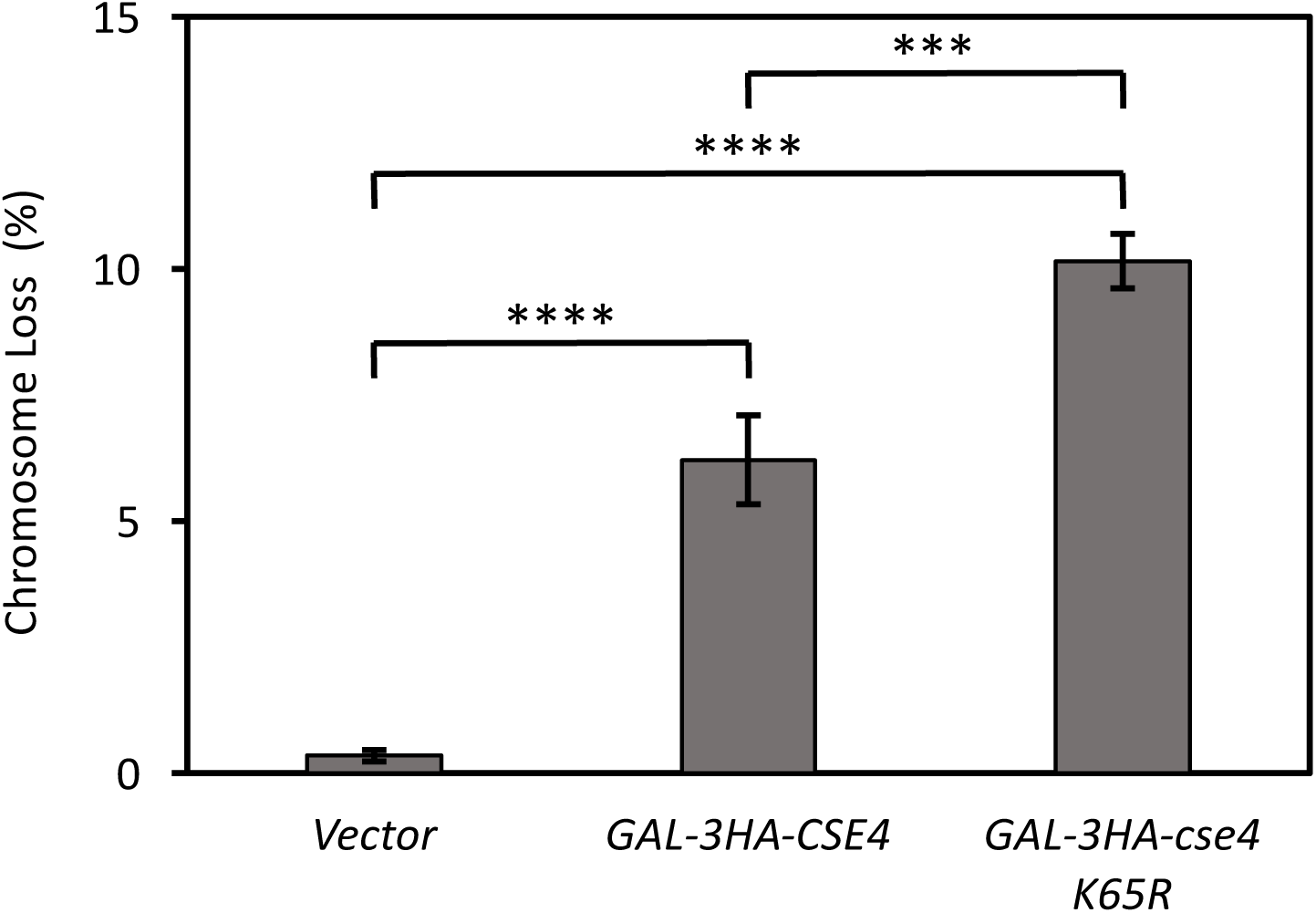
*GAL-cse4 K65R* strains exhibit increased chromosome loss. (A) YPH1018 transformed with vector (pMB433), *GAL-3HA-CSE4* (pMB1597), or *GAL-3HA-cse4 K65R* (pMB1814) were plated on SC –URA with limiting adenine plus 2 % galactose/raffinose and incubated for 5 days at 30 °C. Chromosome loss was determined by counting the number of colonies that show loss of the reporter chromosome in the first division and exhibit a half sector phenotype. Quantification of three independent transformants is shown. Statistical significance was assessed by one-way ANOVA (P < 0.0001) followed by all possible pairwise comparisons of the means with Bonferroni correction. ***, P < 0.001; ****, P < 0.0001.

## Discussion

We have identified Cse4 K65 as a sumoylation site and showed that Cse4 sumoylation regulates its ubiquitin-mediated proteolysis and localization for genome stability. Strains expressing *cse4 K65R* exhibit defects in Cse4 sumoylation and ubiquitination and show reduced interaction of cse4 K65R with Slx5. Defects in proteolysis and mislocalization of cse4 K65R to non-centromeric chromatin were observed under normal physiological conditions. The increased stability of cse4 K65R in *psh1∆* strains but not in *slx5∆* strains indicates that sumoylation of Cse4 K65 regulates Slx5-mediated proteolysis, independently of Psh1. We propose that sumoylation of Cse4 K65 promotes its interaction with Slx5, and this is required for Slx5-mediated proteolysis of Cse4 to prevent its mislocalization for faithful chromosome segregation.

Reduced Cse4 sumoylation observed for the *cse4 K65R* mutant supports the role of Cse4 K65 as a sumoylation site in the N-terminus of Cse4. The defects in sumoylation and ubiquitination of cse4 K65R are not as severe as that observed for cse4 16KR. Furthermore, ubiquitination defects of Cse4 are more pronounced in *slx5∆* strains (Ohkuni et al. 2016), compared to that observed for cse4 K65R. These results suggest that sumoylation and ubiquitination of other lysine residues of Cse4 or other substrates of Slx5 also contribute to proteolysis of Cse4.

Our results show that SUMO modification of Cse4 K65 is required for Slx5-mediated proteolysis independent of Psh1. Several experimental evidences support this conclusion, for example, we observed: 1) reduced interaction of cse4 K65R with Slx5, 2) no additional proteolysis defects of cse4 K65R in *slx5∆* strains, and 3) increased proteolysis defects of cse4 K65R in *psh1∆* strains. Our results are consistent with previous studies which showed ubiquitination of K131, K155, K163 and K172, but not K65 of Cse4, by Psh1 (Hewawasam et al. 2010).

Studies to date have described multiple E3 ligases including Psh1, Slx5, Ubr1, and Rcy1 (F-box protein in the SCF (Skp1-Cullin-F-box) complex) in the proteolysis of overexpressed Cse4 (Cheng et al. 2017). We note that the physiological consequence of defects in the Cse4 E3 ligases characterized to date is most apparent when Cse4 is overexpressed. We propose that this type of regulation ensures that levels of endogenously expressed Cse4 are maintained at a critical threshold for the essential role of Cse4 at centromeres.

In summary, our studies have identified and characterized the physiological role of Cse4 sumoylation and shown that sumoylation of Cse4 K65 regulates ubiquitin-mediated proteolysis by Slx5 to prevent mislocalization of Cse4. Sumoylation of kinetochore proteins or proteins required for sister chromatid cohesion and spindle assembly checkpoint regulate faithful chromosome segregation (Eifler and Vertegaal 2015; Mukhopadhyay and Dasso 2017). Deregulation of sumoylation machinery causes severe defects in cell proliferation and genome stability and promotes tumorigenesis (Eifler and Vertegaal 2015; Seeler and Dejean 2017). Unlike Psh1, Slx5 is evolutionarily conserved and the mammalian orthologue RNF4 has been shown to regulate chromosome segregation (van de Pasch et al. 2013). Since ubiquitination and sumoylation prevent mislocalization of Cse4 for genome stability, it will be of interest to determine if these post-translational modifications of CENP-A also prevent its mislocalization to preserve genome stability in human cells.

## Acknowledgments

We would like to thank members of the Basrai laboratory for helpful discussions and comments on the manuscript. We gratefully acknowledge Mary Dasso and Oliver Kerscher for useful discussions. This work was supported by the National Institutes of Health Intramural Research Program to MAB.

Author contributions: K.O. designed the study, conducted all the experiments, analyzed the data, and wrote the manuscript. R.L.-M. contributed to sumoylation assays, J. W. performed some of the chromosome spreads experiments. W.C.A. generated plasmids and provided technical advice for biochemical assays. Y.T. made comments on the manuscript. R.E.B. performed statistical analyses and assisted with the writing of the manuscript. M.A.B. guided the project and contributed to the writing of the manuscript.

